# Conversion of CO_2_ into valuable products: Engineering the PirC-PGAM switch in cyanobacteria to direct carbon flux into desired products

**DOI:** 10.64898/2026.02.05.703947

**Authors:** Nathalie Sofie Becker, Franziska Hufnagel, Paul Bolay, Kevin Otec, Tim Orthwein, Andreas Kulik, Phillipp Fink, Claudius Lenz, Pia Lindberg, Karl Forchhammer, Stephan Klähn

## Abstract

**Background:** In response to rising CO_2_ emissions driving global warming, there is an urgent need for a transition toward a sustainable bioeconomy. Photo-biotechnological processes based on oxygenic photosynthesis hold high potential for achieving CO_2_ neutrality and in this regard, cyanobacteria have emerged as promising biocatalysts. Rational metabolic engineering of cyanobacteria depends on a thorough understanding of native regulatory mechanisms governing primary metabolism, which can limit metabolic flux through specific pathways and, consequently, the formation of target products. Recent insights have identified a key regulatory node at the 2,3-bisphosphogylcerate-independent phosphoglycerate mutase (PGAM) reaction, where the metabolic flux from newly fixed carbon is redirected from the Calvin-Benson-Bassham (CBB) cycle towards lower glycolysis. This metabolic valve is controlled by the small inhibitor protein PirC, whose binding to PGAM is determined by the central signal transduction protein PII.

**Results:** In this study, we exploit the PirC-PGAM interaction as a novel target for regulatory metabolic engineering in the model cyanobacterium *Synechocystis* sp. PCC 6803 (*Synechocystis*). Chassis strains with engineered control of PGAM, defined as PGAM-ON or PGAM-OFF states, were generated using two complementary approaches: tuning *pgam* gene expression and modulating PirC abundance to regulate PGAM activity. The effectiveness of this regulatory engineering strategy was demonstrated by redirecting carbon flux toward two representative, naturally occurring products: sucrose, produced via gluconeogenesis fueled by the Calvin-Benson-Bassham (CBB) cycle, and succinate, an intermediate of the tricarboxylic acid (TCA) cycle. Narrowing the PGAM valve resulted in a threefold increase in sucrose accumulation. In contrast, opening the PGAM valve by relieving PGAM inhibition through *pirC* deletion or separate *pgam* overexpression resulted in up to an 18-fold increase in succinate excretion. Furthermore, similar genetic configurations were applied to enhance production of a heterologous compound, isoprene, derived from pyruvate.

**Conclusions:** This study establishes the PGAM valve as a tunable control point for the rational re-direction of carbon flux in *Synechocystis* and highlights small regulatory proteins as powerful targets for metabolic engineering. Together, these findings provide proof of concept for an advanced level of molecular engineering in cyanobacteria and to fully harness their biocatalytic potential in future photosynthesis-driven biotechnological applications.

## Background

The anthropogenic climate change leads to a growing need for sustainable, i.e. CO_2_-neutral, biotechnological processes that may also consider photoautotrophic microorganisms as potential production strains. Cyanobacteria are the only prokaryotes performing oxygenic photosynthesis, which makes use of solar energy to oxidize water and utilize the obtained electrons to assimilate CO_2_ [1]. With this, cyanobacteria are dominant primary producers and hence, key players in global biogeochemical cycles [2]. In addition to their environmental impact, they receive growing interest as biocatalysts in photo-biotechnological applications nowadays, e.g. for the sustainable production of value chemicals and fuels [3–5]. To rationally engineer cyanobacteria, e.g. by channeling metabolic fluxes to obtain the maximum yield of a desired chemical product, it is of paramount importance to fully comprehend underlying molecular processes that control primary carbon metabolism [6,7]. Although we are only at the beginning of a full understanding of the regulation of cyanobacterial metabolism, recent research has opened the window into a more comprehensive view on fundamental control principles.

Maintaining the cellular carbon/nitrogen (C/N) homeostasis is one of the most fundamental aspects of cellular physiology. During vegetative growth of autotrophic organisms, it is essential to tightly interconnect CO_2_ fixation with N assimilation. To fulfil this task, cyanobacteria use a sophisticated signaling network centered around the pervasive PII sensor protein [8,9]. PII proteins are fundamental for most free-living prokaryotes and plant chloroplasts, where they act as multitasking signal integrators that combine information on the metabolic C/N balance with the energy state of the cell. C/N sensing is achieved through interaction with the metabolite 2-oxoglutarate (2-OG) and energy sensing by competitive binding of ATP or ADP [9]. 2-OG is ideally suited as a status reporter for C/N balance sensing, as it is the C skeleton, into which ammonium is incorporated through the assimilatory reactions catalyzed by the glutamine synthetase-glutamate synthase (GS/GOGAT) cycle [10].

The mechanistic details, how the PII proteins interact with effector molecules and transmit this information into distinct conformational states, have been elaborated in great detail [9,11]. Briefly, differential binding of the effector molecules ATP/ADP and 2-OG to the PII protein affects the conformation of flexible loop regions of PII, and depending on these conformational states, PII interacts with its receptor proteins, and by that modulating their activities. By this mechanism, PII controls key enzymes, transporters and small regulatory proteins. For example, PII controls the transcriptional co-activator protein PipX, which the global nitrogen control factor A (NtcA) requires to activate the expression of various N assimilatory genes [12,13]. Furthermore, PII controls the activity of enzymes directly, e.g. acetyl-CoA carboxylase or N-acetyl-L-glutamate-kinase (NAGK), which catalyzes the committed step in arginine synthesis [14–16]. More recently, interactions of PII with small proteins of previously unknown function were also revealed, which in turn provided a plethora of new insights (see below).

The interest in small proteins (< 100 amino acids) and their important role in cellular functions has only recently gained more attention [17]. Cyanobacteria appear to provide a paradigm in this regard, as the photosynthetic apparatus contains numerous proteins of less than 50 amino acids [18]. Moreover, several small proteins act as effectors of key enzymes [19] and PII interacts with several of these small proteins. For example, the small adaptor protein PipX, a co-activator of the global transcriptional regulator NtcA, is bound to PII in N replete conditions and is released upon N limitation to form a complex with NtcA bound to 2-OG, thereby co-activating NtcA [20]. Furthermore, we showed recently the 51 amino acid protein PirA to antagonize functions of PII by binding PII in an ADP-dependent manner and in turn preventing activation of N-acetylglutamate kinase [21].

The 2,3-bisphosphoglycerate-independent phosphoglycerate-mutase (PGAM) reaction, which converts the first CO_2_ fixation product 3-phosphoglycerate (3-PGA) to 2-phosphoglycerate (2-PGA), represents another major control point of the C/N balance that also involves the PII protein [22]. The activity of PGAM determines the route of newly fixed carbon. In particular, C is taken out from the CBB cycle and redirected towards lower glycolysis, supplying the carbon skeleton for important anabolic reactions in amino acid, nucleic acid and lipid metabolism. PGAM activity is controlled by the interaction with the 112 amino acid protein PirC (**P**II-**i**nteracting **r**egulator of **c**arbon metabolism; *sll0944* product, which has also been termed CfrA [23]). PirC itself is under control of the PII protein [22,24]. Under N replete conditions, hence low 2-OG levels, PirC is sequestered by PII and inactive. When the N supply becomes limiting, the cellular 2-OG levels increase, leading to the PII-ATP-2OG conformation, which releases PirC from the PII protein complex. In turn, free PirC binds to and inhibits the activity of PGAM, thereby slowing the conversion of 3-PGA to 2-PGA, which triggers the rapid synthesis of glycogen through activation of GlgC by increasing 3-PGA levels [25]. Similarly, overexpression of the *pirC* gene caused rapid accumulation of glycogen under vegetative conditions [23]. These results highlight the PirC-PGAM switch as a pivot point for controlling the carbon flow in cyanobacterial metabolism.

Compared to classical metabolic engineering, focusing on the introduction of novel pathways, targeting regulators for the intracellular re-modulation of metabolic fluxes is less prevalent but also emerging in cyanobacteria [6,7,26]. Here, we engineered the cyanobacterial model strain *Synechocystis* sp. PCC 6803 by tuning the PirC-PGAM switch and provide a proof of concept, by modulating the components PGAM and PirC, to reach the maximum flux either into the direction of acetyl-CoA (PGAM valve open, i.e. PGAM-ON state) or into gluconeogenic direction (PGAM valve closed, i.e. PGAM-OFF state). The PGAM-ON state was achieved via overexpression of the *pgam* gene, deleting the *pirC* gene or a combination of both. Under conditions of nitrogen starvation, the PGAM-ON state led to excretion of metabolites from lower glycolysis, such as pyruvate, 2-OG and succinate and to increased accumulation of PHB, proving the redirection of carbon-flow towards lower glycolysis. Moreover, a PGAM-ON state in a recombinant strain also expressing a plant-derived isoprene synthase led to increased production of isoprene under N deprivation. Conversely, a PGAM-OFF state was established through inducible *pirC* overexpression combined with *pgam* downregulation. Carbon flux redirection was evidenced by increased glycogen accumulation, and optimized configurations were applied to enhance sucrose production in salt-stressed *Synechocystis*. After induction, PGAM-OFF strains accumulated more glycogen compared to the wild type, and also showed elevated sucrose levels under high-salt conditions.

## Materials and Methods

### Strains and growth conditions

*Synechocystis* sp. PCC 6803 was obtained from the Pasteur Culture Collection of Cyanobacteria and used as the wild type strain. Cultivation was carried out in BG11 medium [27]. Experimental cultures were adjusted to an OD_750_ of 0.4 and maintained at 28°C under continuous illumination (∼70–80 µmol photons m⁻² s⁻¹), with shaking at 125 rpm. For BG11 agar plates, the medium was supplemented with 20 mM sodium tris-(hydroxymethyl)-ethyl phosphonate buffer (pH 8.2) and 1.5% Bacto agar. When required, appropriate antibiotics were added to pre-cultures. For nitrogen depletion experiments, cells were pre-grown in BG11 medium to an OD_750_ of 0.6–1.0, then washed three times with nitrogen-free BG11 medium (BG11^0^). For glycogen and sucrose measurement during PGAM-OFF state, cultures were inoculated at an OD_750_ of 0.2 and cultivated in Cu^2+^-free BG11 with appropriate antibiotics. For induction of the PGAM-OFF state CuSO_4_ was added to a final concentration of 1 µM. The cultures were then cultivated at 30 °C, 150 rpm, ambient CO_2_, 50 µmol photons m^-2^ s^-1^ and 70% relative humidity. *Escherichia coli* (*E. coli*) strains were cultured in LB medium or on LB agar plates (10 g/L NaCl, 10 g/L tryptone, 5 g/L yeast extract) at 37 °C, supplemented with the appropriate antibiotic when required.

### Genetic engineering

All plasmids used and generated in this study are given in **Table S1**, the obtained recombinant strains are listed in **Table S2**. The oligonucleotides used for DNA amplification are given in **Table S3**. To construct the pSEVA251-P*_J23101_*::*pgam* plasmid, Gibson Assembly was used [28]. DNA amplification was performed with Q5 High-Fidelity DNA Polymerase (NEB) (primer NB042/045/046/047) using the pSEVA251 plasmid or genomic *Synechocystis* DNA as template. For purification of PCR-products and plasmid isolation, the Monarch^®^ DNA Gel Extraction Kit and Monarch^®^ Plasmid Miniprep Kit (NEB) were used, respectively. The pSEVA251-P*_J23119_*::*pgam* plasmid was constructed using the Q5 Site-Directed Mutagenesis Kit (NEB) (primer NB075/076). Plasmids were checked via sequencing (Eurofins Genomics, Ebersberg, Germany). To construct the pSEVA451-P*_petE_::pirC* plasmid a restriction-ligation approach was used. For this, the respective promoter and gene sequences were amplified from the *Synechocystis* gDNA with primers adding the recognition sites of the restriction enzymes used (FH013/017/018/019) and fused by PCR (FH013/019). The fused gene construct was introduced into a pJET plasmid (CloneJET PCR Cloning Kit, Thermo Fisher Scientific, Darmstadt) and excised from it using *Eco*RI and *Bam*HI. The gene construct was then ligated into a pSEVA451 plasmid digested with the same enzymes using T4 ligase (Thermo Fisher Scientific, Darmstadt). For purification of PCR-products and plasmid isolation the NucleoSpin Gel and PCR clean-up Kit (Macherey-Nagel, Düren) and the NucleoSpin Plasmid Mini Kit (Macherey-Nagel, Düren) were used, respectively. The pEX-K248-*P_petJ__pgam* plasmid was synthesized by Eurofins. Plasmids were checked via sequencing (Genewiz, Azenta Life Sciences, South Plainfield NJ).

The plasmids pSEVA251-P*_J23101_*::*pgam* and pSEVA451-P*_petE_::pirC* were introduced into *Synechocystis* via electroporation as described [29], with minor modifications. To make *Synechocystis* electrocompetent, 30 mL of culture in linear phase around OD_750_ 0.5 was collected (4,000 g, 15 min, 4 °C). Cells were washed 3 times with 20 mL ice-cold HEPES (4-(2-hydroxyethyl)-1-piperazineethanesulfonic acid) buffer (1 mM, pH 7.5). Then, cells were resuspended in 1 mL HEPES and aliquots of 60 µL were either used directly or stored at -80 °C. Electroporation was done with either 18.0 kV/cm or 2500 V for 5.2 ms, using 100-500 ng plasmid. Cells were resuspended in 800 µL BG11 medium and plated onto BG11 agar plate without antibiotics and incubated at 28 °C for one night before increasing the antibiotic concentration over the next days. Clones were validated with colony PCR (primer NB062/063 or primer FH017/019/076/081).

The *P_petJ_::pgam* configuration was achieved by homologous recombination applying the pEX-K248-*P_petJ__pgam* plasmid on naturally competent *Synechocystis*. For this, WT cells were collected in their logarithmic growth phase at an OD_750_ of 0.5 to 1 by centrifugation at 3,000 x g for 5 min. The supernatant was discarded and the cells were resuspended in BG11 (in 1% volume of initially sampled material). Around 100 to 200 ng of DNA were added to 100 µL of concentrated cells, mixed gently and incubated overnight at 30 °C in the dark and shaken at 150 rpm. Cells were plated on BG11 with a low concentration of antibiotics and cultivated at 30 °C. Once colonies appeared, they were picked onto plates with an increased concentration of antibiotics. Clones of the *pgam_KD* strain were validated with colony PCR (FH011/020/027/101/102/111).

For the generation of isoprene producing *pirC* deletion strains, the isoprene production cassette was amplified from plasmid pEERM3-2MEP-IspS:KmR via primers 2-MEP-PirCup_fw/rev. The plasmid backbone was amplified with primers pEERM-PirCdwn_fw/pEERM-PirCup_rev. 500 bp homology regions up- and downstream of *pirC* were generated with primers PirCup_fw/PirCup_2MEP_rev and PirCwnd-2MEP_fw/PirCdwn_rev, respectively. The resulting constructs were fused via AQUA cloning [30] and analogously used to obtain recombinant *Synechocystis* strains.

### Immunoblotting

To analyze relative PGAM and PirC levels in recombinant *Synechocystis* strains, cells from 5-10 mL culture were harvested by centrifugation at 4,000 x g for 15 min and the cell pellet was stored at -20 °C until use. For cell lysis, the pellet was resuspended in 100 µL extraction buffer (50 mM Tris-HCl, 5 mM EDTA, 100 µg/mL lysozyme (from chicken egg white), 1 µL/100 µL protease inhibitor (Sigma Aldrich)). The samples were then incubated on ice for 1 hour. An equal volume of glass beads (0.1-0.15 mm) was added for cell disruption using a FastPrep-24 Ribolyser (MP Biomedicals) with five cycles of 15 s at 6.5 m/s and 4 °C. To load protein extracts on the SDS-PAGE that correspond to equal volume of culture per lane, the samples were normalized to the autofluorescence. This was done to correct different cell lysis efficiency, as well as drastic changes in protein amounts in N-starved cells. The fluorescence of the cultures was measured immediately before the cell harvest by a WATER-PAM chlorophyll fluorometer (Walz GmbH, Effeltrich, Germany). The autofluorescence of the culture at time 0 was set in relation to the autofluorescence measured at each time point of nitrogen-depletion. After lysis, the autofluorescence was measured in the lysates and the same ratio as between the cultures was adjusted by correcting the volume size with the following correction factor (see formula 1). For the control sample at time point 0, ∼10 µg protein per well were loaded, while for the samples in N starvation the correction factor was used to calculate the volume accordingly.

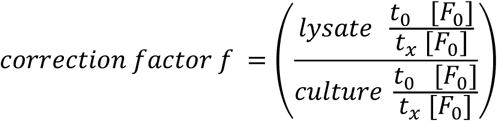

**Formula 1: Determination of the correction factor for SDS-PAGE sampling**. To determine the volume of the SDS-PAGE sample, a correction factor (f) was determined using the ratio of the measured base fluorescence (F_0_) in the lysate normalized to the ratio of the fluorescence in the cell culture. The ratio thereby represents the fluorescence at a time point x in chlorosis (t_x_) to the control time point (before chlorosis, t_0_). F_0_ = base fluorescence; t_0_ = control time point (before chlorosis), t_x_ = time point x in chlorosis.

After measuring the autofluorescence of the lysates, the samples were centrifuged (16,000 g, 5 min, 4 °C) and the calculated volumes of lysate were boiled with SDS-loading dye (95 °C, 10 min). SDS-PAGE was performed as previously described [31], using 4-15% precast midi gels (Q-PAGE). Western blotting was performed as previously described [32], using a methanol-activated PVDF membrane (ROTI^®^PVDF 0.45). After protein transfer, the membrane was blocked with 5% (w/v) milk powder in TBS-T buffer (50 mM Tris, 150 mM NaCl, 0.1% Tween-20, pH 7.4). After washing with TBS-T buffer, the membrane was incubated with the primary antibody (PGAM or PirC antiserum from rabbit, Davids Biotechnologie (see Appendix), 1:3,000 dilution in TBS-T) for 1-2 h at RT, followed by a wash step. Then, the secondary antibody (antirabbit polyclonal goat antibody–horseradish peroxidase conjugate; Sigma-Aldrich, 1:5,000 dilution in TBS-T) was incubated for 1-2 h at RT or at 4 °C overnight, followed by a wash step. Immunoreactive bands on the membrane were visualized using the LumiLight detection system (Roche Diagnostics) and the Gel Logic 1500 imaging system (Kodak) with the respective software.

### Metabolite profiling

#### Glycogen quantification

Method A: Glycogen content was quantified using a colorimetric assay with *o*-dianisidine as recently described [33]. Method B: Glycogen was additionally quantified by another colorimetric method using *o*-toluidine adapted from [34]. For this, 2 mL of the cultures were centrifuged at 16,000 x g for 10 min. The pellet was washed twice with 1 mL MiliQ-water and suspended in 400 µL of KOH (30% (w/v)) and incubated at 95 °C for 2 h. Afterwards, 1.2 mL of ice-cold ethanol (abs.) was added and glycogen was precipitated at -20 °C overnight. The samples were centrifuged at 10,000 x g for 10 min and the supernatant was discarded. The cells were first washed with 500 µL ethanol (70%) and then with 500 µL ethanol (abs.). The pellet was dried for 20 min at 60 °C in the Speed-Vac (Concentrator plus/Vacfuge plus®, Eppendorf, Wesseling-Berzdorf). The dried pellet was taken up in 1 mL of 100 mM sodium acetate and 8 µL of amyloglucosidase (4.4 U/µL) were added. The samples were incubated for 2 h at 60 °C. 200 µL of the sample were mixed with 1 mL *o*-toluidine-reagent (6% in glacial acetic acid (v/v)) and incubated at 100 °C for 10 min. Afterwards, the reaction was stopped by incubation on ice for 3 min. The measurement was done with 100 µL in a 96-well plate at 635 nm in a microwell-reader (Infinite 200 PRO Reader, Tecan, Männedorf). As a standard, different glucose concentrations (1 mg/mL, 500 µg/mL, 250 µg/mL, 125 µg/mL, 62.5 µg/mL, 31.25 µg/mL and 0 µg/mL), also incubated with *o*-toulidine-reagent as described above, were used.

#### PHB quantification

PHB was quantified with HPLC as described before [35], with the minor change that 10 mL cell cultures were harvested and cell pellets were dried at 60 °C until the supernatant evaporated.

#### Sucrose quantification

For sucrose quantification, 2 mL cultures were centrifuged at 16,000 x g for 10 min and the cell pellet was stored at -20 °C. Sucrose was extracted adapting a strategy from literature [36]. The pellet was taken up in pre-cooled 80% ethanol. The cells were transferred into lysis tubes with glass beads (Precellys® Aufschluss-Kit, Bertin technologies, Montigny-le-Bretonneux) and lysed using a homogenizer (Precellys® evolution, Bertin technologies, Montigny-le-Bretonneux) and afterwards incubated at 55 °C for 10 min. The suspension was centrifuged at 12,000 x g for 45 min and transferred to a fresh 2 mL tube. The ethanol was evaporated in a Speed-Vac (Concentrator plus/Vacfuge plus®, Eppendorf, Wesseling-Berzdorf) at 30 °C for 2 h. The pellet was taken up in 200 µL of MiliQ water. Sucrose was measured using a Vanquish HPLC system (Thermo Scientific, Waltham) system equipped with an HiPlex H column (300 mm x 7.7mm x 8µm, Agilent, Santa Clara) and a refractive index detector. Water at a flowrate of 0.6 ml/min was used as the eluent. Column temperature was 15°C. As a standard, sucrose (Applichem, Darmstadt) at 1 mM, 0.5 mM, 0.25 mM, 0.125 mM, 0.0625mM and 0.03125 mM in MilliQ water was used.

#### Quantification of extracellular metabolites

Extracellular pyruvate, succinate, 2-oxoglutarate and glutamic acid concentrations were analyzed with HPLC-MS from cell-free supernatant as described previously [33]. As standard, 0.01, 0.05 and 0.1 mM of sodium pyruvate (C_3_H_3_NaO_3_, Sigma-Aldrich), sodium succinate dibasic hexahydrate (NaOOCCH_2_CH_2_COONa · 6 H_2_O, Sigma-Aldrich) and α-ketoglutaric acid disodium salt hydrate (2-OG, C_5_H_4_O_5_Na_2_· xH_2_O, Fluka Analytical) were used.

#### Assessment of isoprene production

For determining isoprene production, 20 mL of seed culture without antibiotics was transferred to 60 mL air-tight vials (Fisherbrand cat. no. 11540585) and cultivated at 40°C, 80 µmol photons m^-2^ s^-1^ and 150 rpm for 1-15 h. For isoprene quantification, 150 µL of headspace were injected in a gas chromatograph (Clarus 580, Perkin Elmer, Waltham, USA) equipped with a flame-ionisation detector and a Porapak QS packed column (Porapak QS 80/80 PE 8000, 1.8 m × 2 mm ID, Cat. No. N9305013-ZW5531, Perkin Elmer). Oven temperature was set to 200°C for 2.5 min with N_2_ at 20 mL min^−1^ as carrier gas. Isoprene elution peaked at 1.74 min and was quantified based on the peak area and synthetic standards. OD_750_ was determined after sampling.

## Results

To demonstrate the potential of regulatory engineering to re-direct carbon-flow towards desired product formation in cyanobacteria, we have generated *Synechocystis* strains with modified PGAM valves. Two different directions of carbon-flow have been considered. For the synthesis of products derived from lower glycolysis, the PGAM valve must be maximally open (PGAM-ON). To synthesize products, derived from the upper part of central carbon metabolism (sugars, glycerol), the PGAM valve has to be completely closed (PGAM-OFF). To achieve these states, we modulated the abundance of either PGAM or PirC (**Fig. 1**).

**Figure 1:**
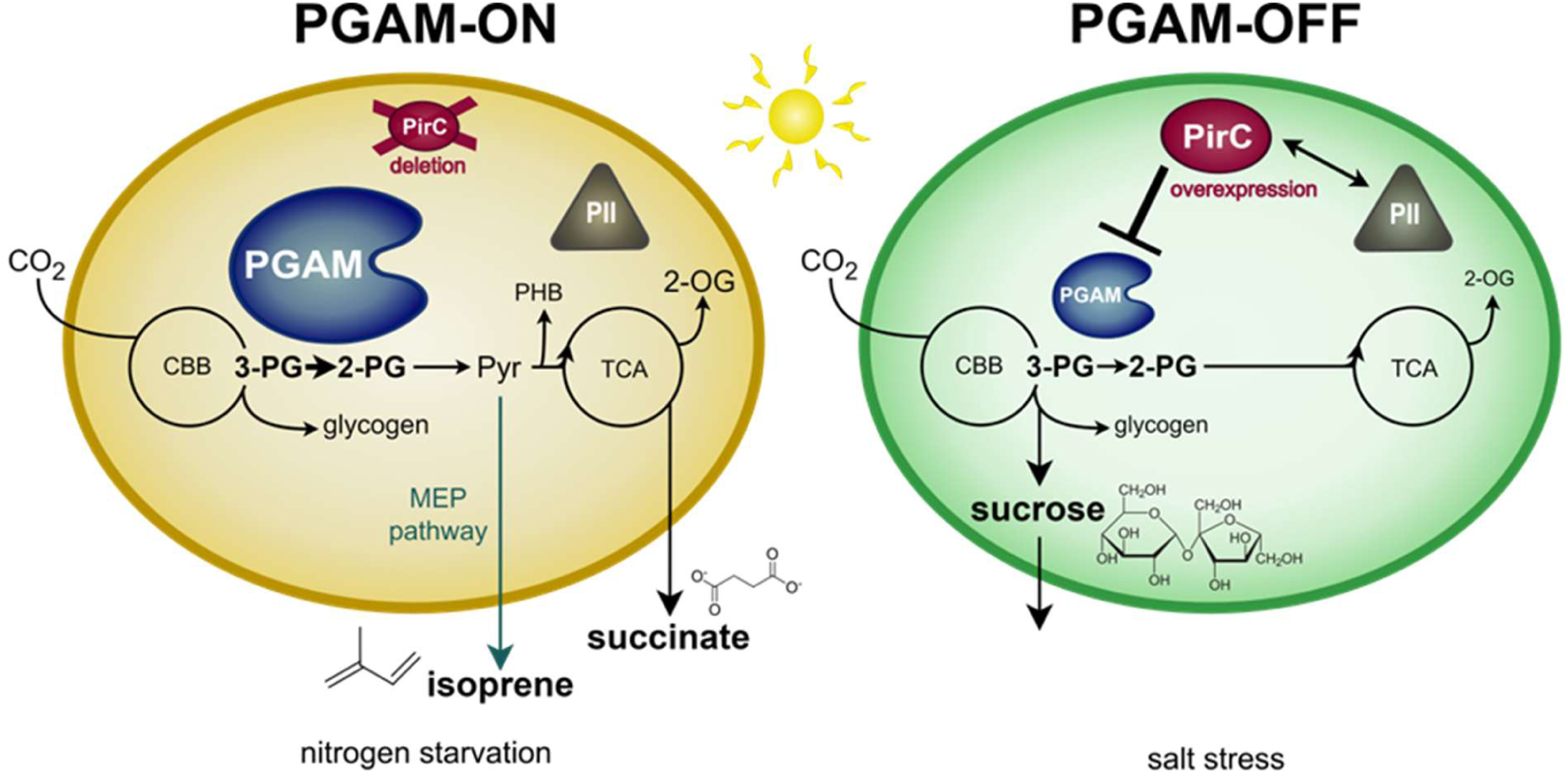
Illustration of the basic principle of engineering the PirC-PGAM-valve. In *Synechocystis*, the PGAM-PirC switch tightly regulates the carbon flow at the branching point between upper and lower glycolysis. In this case study, the aim is to direct the carbon flow either towards lower glycolysis (succinate and isoprene excretion), by overexpressing PGAM to decrease the inhibition of PirC under nitrogen starvation (PGAM-ON state, left panel) or towards gluconeogenesis (sucrose production), by engineering a strong inhibition of PGAM by PirC (PGAM-OFF state, right panel).

### Regulatory engineering towards the PGAM-ON state in *Synechocystis*

A potential PGAM-ON state requires high basal PGAM activity, which might be obtained by increasing enzyme abundance via overexpression of the *pgam* gene. To achieve the PGAM-ON state, *Synechocystis* strains were generated that ectopically express the *pgam* gene driven by constitutive synthetic promoters (BioBricks BBa), either J23101 for moderate or J23119 for strong overexpression, similar to previous studies [37,38] (**Fig. 2A**). These plasmids were introduced into *Synechocystis* WT and into a strain lacking the *pirC* gene [22], which therefore lacks the inhibition of PGAM by PirC. Finally, we obtained the strains 101_*pgam*, 119_*pgam* and Δ*pirC* + 101_*pgam*, which were verified by colony PCR (see **Fig. S1A-C**), and we performed immunoblotting using a PGAM-specific antiserum. As expected, when *pgam* gene expression was driven by either PJ23101 or PJ23119, strong PGAM accumulation was achieved compared to the WT or Δ*pirC* control strains (**Fig. 2B**). In addition to vegetative growth conditions, we also analyzed cells grown under nitrogen starvation as *pgam* gene expression was reported to be downregulated in nitrogen-starved cells, according to the microarray analysis shown in [39]. However, PGAM overproduction in the engineered strains was not affected by nitrogen starvation and remained stably high even after 14 d in nitrogen starvation (**Fig. S1D**). Under either condition, the expression driven by the J23119 promoter led to higher PGAM abundance compared to the J23101 promoter, which is in agreement with previously reported promoter strength [37]. Contrary to its interaction target, PirC is increased under nitrogen starvation (**Fig. 2B**), which is in agreement with previous reports [39,40]. Interestingly, we observed a correlation between increased PGAM abundance in strains 101_*pgam* and 119_*pgam* and an increase of PirC (**Fig. 2B**) even though the *pirC* gene expression was not genetically manipulated in these strains. The PGAM-dependent accumulation of PirC might be a response to counteract a redirection of carbon flow towards lower glycolysis albeit the mechanism behind this remains unclear.

**Figure 2:**
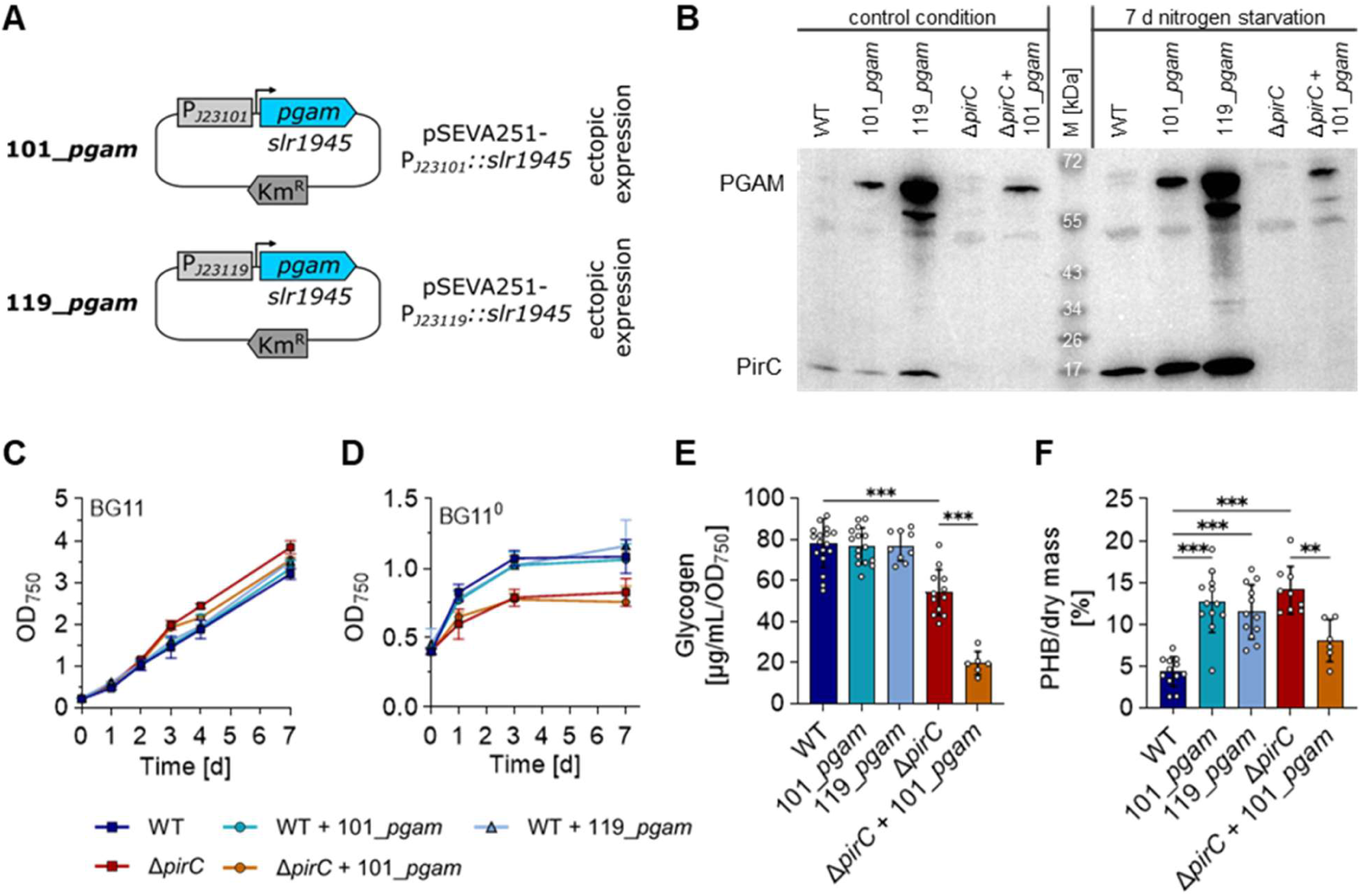
Demonstration of the PGAM-ON state. **A:** Plasmid design for overexpressing the native *pgam* gene (*slr1945*) from the pSEVA251 plasmid under control of the moderate J23101 or strong J23119 promoter. **B:** Immunoblots against PGAM and PirC for cells grown under vegetative (control) conditions and for 7 days under N starvation. **C-D:** Vegetative growth (C) and OD_750_ under nitrogen starvation (D) with at least three biological replicates. **E-F:** Quantification of glycogen with method A (E) and PHB (F) after 7 days of N starvation. Each dot represents a biological replicate. Dunn’s multiple comparisons test was used (ns = P > 0.05, * = P ≤ 0.05, ** = P ≤ 0.01, *** = P≤ 0.001).

After detecting higher PGAM levels in the *pgam* overexpressing strains, we analyzed if the established genetic configurations might affect growth performance of *Synechocystis*. As compared to the corresponding reference strains, neither the presence of pSEVA251 nor the overexpression of *pgam* had a significant impact on growth performance under vegetative or nitrogen-depleted conditions (**Fig. 2C, D** & **Fig. S2**). As already reported previously, upon nitrogen-downshift the Δ*pirC* mutant didn’t show the final doubling before arresting growth that *Synechocystis* usually does, but showed a premature growth arrest [22,24]. The Δ*pirC* + 101_*pgam* strain behaved in the same way as the Δ*pirC* strain.

Previous studies showed that a Δ*pirC* mutant, i.e. a strain lacking the proteinaceous inhibitor of PGAM, produces less glycogen and more PHB under nitrogen starvation compared to the WT [22]. However, under this uninhibited condition, the amount of PGAM might still be the limiting factor of carbon flow. To evaluate this, we made a similar experiment as Orthwein et al. (2021) using the generated *pgam* overexpression strains and analyzed glycogen and PHB contents after 7 days of N starvation (**Fig. 2E, F**). Indeed, the *pgam* overexpression altered the carbon flow, but with different outcome in either WT or Δ*pirC* background. In the WT background, glycogen content was not significantly altered by higher PGAM levels, neither in the 101_*pgam* nor in the 119_*pgam* strain (**Fig. 3E**). However, both PGAM overproduction strains produced around 12% PHB per dry mass, which was three times more than detected in the WT (∼4%) and the same as in the Δ*pirC* mutant (∼14%) (**Fig. 3F**). In contrast, the Δ*pirC* + 101_*pgam* strain formed significantly less glycogen compared to the Δ*pirC* reference strain (∼2.5 times less) and PHB production was reduced by almost half (8%). The combined effect of *pirC* deletion and *pgam* overexpression might imbalance cell metabolism more drastically than the *pirC* deletion alone.

**Figure 3:**
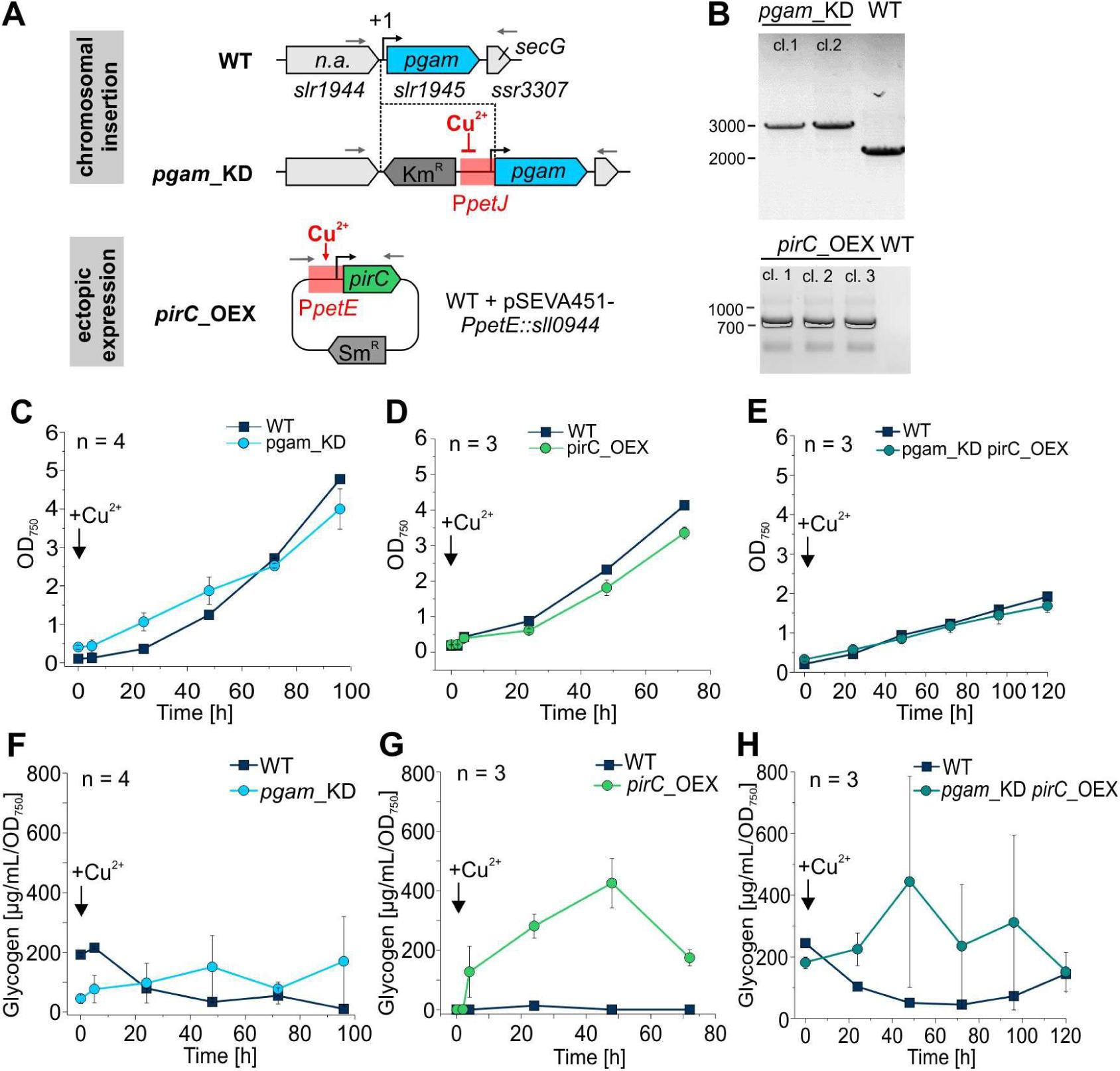
*Synechocystis* strains with genetic configurations demonstrating the PGAM-OFF state. **A:** Design of constructs to achieve *pirC* overexpression and *pgam* downregulation using copper controllable promoters from *Synechocystis.* **B:** Verification of recombinant strains. The obtained strains were verified by PCR, using primer combinations illustrated by grey arrows in panel A. **C-E:** Growth of the different mutants pgam_KD and pirC_OEX and the combination mutant of pgam_KD and pirC_OEX compared to the wildtype under inducing conditions, i.e in presence of Cu^2+^ at a final concentration of 1 µM. **F-H:** Development of the average glycogen amount in the obtained strains compared to the WT after the addition Cu^2+^ at a final concentration of 1 µM. Glycogen was measured using method B. Data are the mean ± SD of n biological replicates as indicated.

### Regulatory engineering towards the PGAM-OFF state in *Synechocystis*

To achieve the PGAM-OFF state, we aimed at diminishing *pgam* gene expression combined with an overexpression of the *pirC* gene to change the equilibrium towards increased amounts of the PirC-PGAM complex (see **Fig. 1**). This, in turn, would result in inhibited flux towards acetyl-CoA and would boost the yields of products derived from gluconeogenesis, such as sucrose. For this, we fused the corresponding genes with promoters that are controlled by micromolar concentrations of copper ions [41], i.e. the *pirC* gene with the *petE* promoter (P*petE*) and the *pgam* gene with the *petJ* (P*petJ*) promoter, to achieve this scenario upon Cu^2+^ addition (**Fig. 3A**). The obtained construct P*petE*::*pirC* was introduced into *Synechocystis* WT on a replicative plasmid. However, the construct P*petJ*::*pgam* was introduced into the chromosome of *Synechocystis,* thereby replacing the native *pgam* gene locus (**Fig. 3A**). The obtained strains for Cu^2+^-inducible *pgam* knockdown (pgam_KD) or *pirC* overexpression (pirC_OEX) were genetically confirmed (**Fig. 3B**). In addition, a strain that combined both the pgam_KD and the pirC_OEX constructs was designed. Subsequently, all strains were analyzed regarding growth performance to estimate any pleiotropic effects caused by the changed genetic configuration, but none of the configurations did affect growth compared to the WT (**Fig. 3 C-E**). Finally, glycogen accumulation was also tracked in the mutant strains (**Fig. 3 F-H**). Both the pgam_KD and the pirC_OEX strain accumulate glycogen after supplementation with 1 µM copper sulfate, while the glycogen amount within the cultures before induction was comparable to the WT. In particular, strain pgam_KD showed a 6-fold increase of glycogen compared to the WT at 48 h after diminishing *pgam* gene expression (**Fig. 3F**). The effect was more drastic in strain pirC_OEX and levels around 400 µg/mL/OD_750_ at 48 h after induction, while almost no glycogen was detectable in the WT (**Fig. 3G**). The combination of both pgam_KD and pirC_OEX also showed an increased glycogen concentration compared to the WT after addition of Cu^2+^, while the glycogen concentration before the induction with Cu^2+^ was comparable to the WT. However, the effect that could be seen here was not as drastic as the effect of the pirC_OEX on its own and accompanied with rather high biological variation (**Fig. 3H**).

### The PGAM-ON state enhances the production of succinate - a representative of the lower glycolysis route

First of all, the engineered PGAM-ON strains, i.e. those with a modulated carbon flow towards the TCA cycle, were analyzed regarding metabolites known to be excreted from *Synechocystis*. Pyruvate, 2-OG and succinate were reported to be released to the medium under nitrogen depletion, especially when the carbon flow has been altered [33,42,43]. Therefore, extracellular concentrations of these metabolites were quantified in the supernatant of cell cultures that have been N-starved for 7 days using HPLC-MS. Under the conditions used in this study, pyruvate could not be detected in the supernatant of WT cultures under N starvation, similar to previous studies [43]. However, overexpression of *pgam* resulted in increased pyruvate secretion, as in both strains 101_*pgam* and 119_*pgam,* pyruvate was detectable in a range between 2-3 mg/L (**Fig. 4A**). The Δ*pirC* mutant excreted even more pyruvate, which, however, was not further elevated when the *pgam* gene was ectopically expressed. The supernatant of both strains, Δ*pirC* and Δ*pirC*+101_*pgam*, contained ∼4 mg/L pyruvate (**Fig. 4A**). Similar effects were also observed for 2-OG. While ∼1.5 mg/L 2-OG were detected in the medium of WT cultures, strains 101_*pgam* and 119*_pgam* excreted 7-fold and 9-fold higher 2-OG levels compared to the WT, respectively (**Fig. 4B**). Deletion of *pirC* resulted in approximately 20 times higher 2-OG concentrations in the medium, reaching around 28 mg/L, which again did not further increase when *pgam* overexpression was also established.

**Figure 4:**
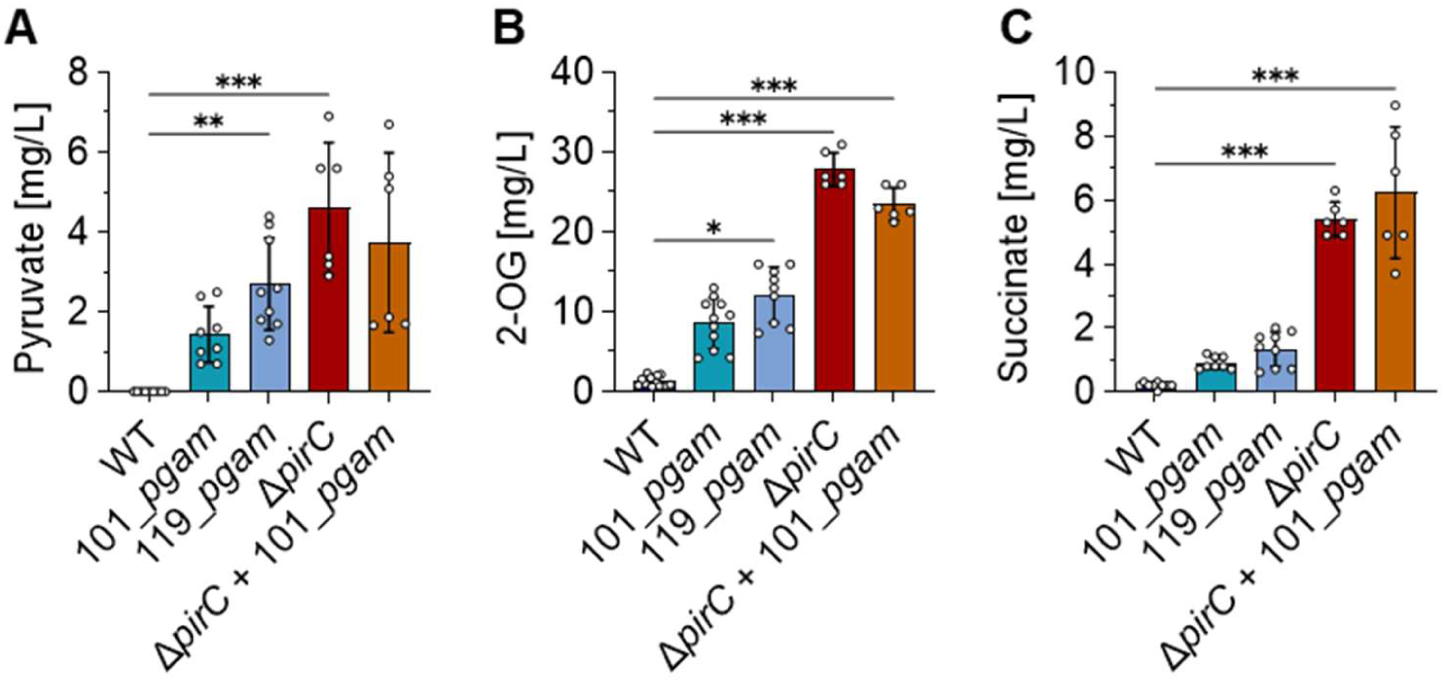
Making use of the PirC-PGAM-switch to form certain products from lower glycolysis, e.g. succinate. Pyruvate (A), 2-OG (B) and succinate (C) were quantified with HPLC-MS from the supernatant of 7-day N-depleted cell cultures. Each dot represents a biological replicate. Dunn’s multiple comparisons test was used (ns = P > 0.05, * = P ≤ 0.05, ** = P ≤ 0.01, *** = P≤ 0.001).

The TCA cycle intermediate succinate was also quantified. The WT strain released 0.3 mg/L succinate to the medium after cultivation for 7 days under N starvation. Again, the established PGAM-ON state resulted in increased succinate excretion compared to the WT (**Fig. 4C**). Strains 101_*pgam* and 119*_pgam* showed comparable succinate levels of between 0.8 and 1.3 mg/L. Accordingly, overexpression of the *pgam* gene resulted in 2- to 4-fold higher succinate excretion. This became much more drastic in the Δ*pirC* strain, which accumulated 5-6 mg/L and hence, 18-fold higher succinate levels compared to the WT (**Fig. 4C**). As seen for the other quantified metabolites, additional *pgam* overexpression in Δ*pirC* could not significantly enhance succinate excretion.

### An opened PGAM valve enhances heterologous production of isoprene

In addition to succinate, which is of biotechnological interest itself [44], we analyzed if the here established genetic configurations, targeting the PGAM valve, also enhance the production of other pyruvate-derived compounds of economic importance. For example, the hydrocarbon isoprene is not naturally produced by *Synechocystis* but can be generated from precursors of the non-mevalonate (MEP) pathway by introducing plant-derived isoprene synthases [45] (**Fig. 5A**). To test the effect of an open PGAM valve on isoprene production, a gene cassette enabling isoprene production was introduced into the *pirC* gene locus via homologous recombination, thereby replacing the native *pirC* gene (**Fig. 5B**). For comparison, a previously published production strain [46] was used, that carries the same isoprene production cassette in ORF *slr0168*, which is dispensable for *Synechocystis* under standard growth conditions and hence considered as neutral integration site (defined as neutral site I (NSI)) (**Fig. 5B**). Here, absence of *pirC* did not have significant effects on growth or isoprene formation under standard conditions (**Fig. 5C, D**), however, 24 hours after induction of N starvation, strong differences in the isoprene output were observed (**Fig. 5E**), pointing towards an open PGAM valve that upholds carbon flux into MEP. Nevertheless, isoprene formation ceased entirely after 3 days, pointing towards complete metabolic arrest under these conditions (not shown). Moreover, volumetric productivity was substantially lower when compared to N-replete conditions, given that growth was arrested entirely, consistent with previous reports.

**Figure 5:**
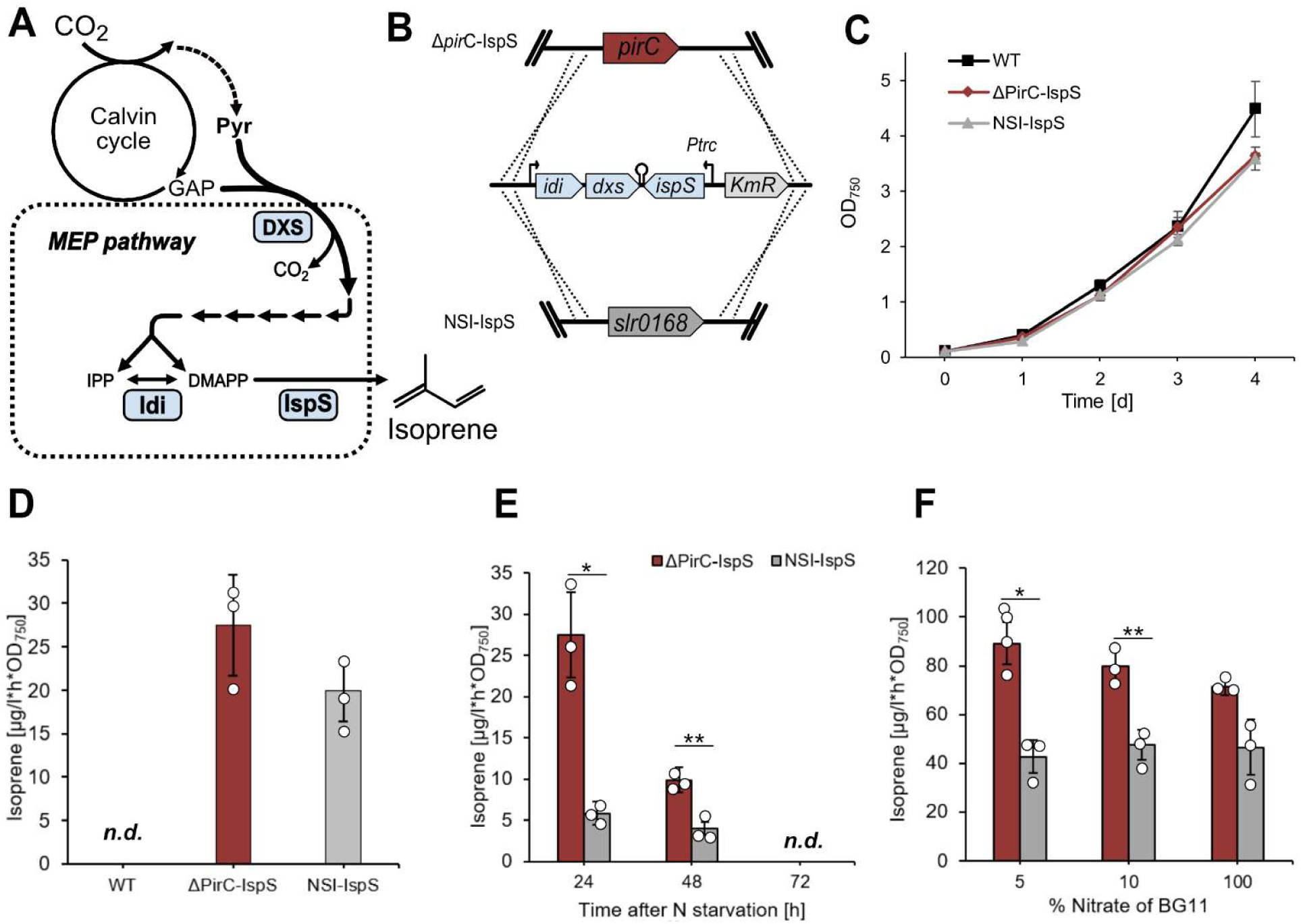
Isoprene formation upon *pirC* deletion in engineered *Synechocystis* strains. **A:** Simplified scheme of metabolic framework for MEP-pathway derived isoprene production, overproduced enzymes in production strains are marked in blue. *Coleus forskohlii* D-xylulose-5-phosphate synthase (DXS), *Synechocystis* isopentenyl pyrophosphate isomerase (Idi), *Eucalyptus globulus* isoprene synthase (IspS) **B:** Genetic constructs enabling isoprene production in *Synechocystis* and concomitant deletion of *pirC* or *slr0168*, respectively. **C:** Growth comparison during standard growth conditions. **D:** Specific isoprene production activity after 3 days of cultivation at standard conditions. **E:** Specific isoprene production activity in response to N starvation. **F:** Specific isoprene production activity after 24 hours of cultivation under respective conditions. Data shown are average values and standard deviations of 3 biological replicates, significance was tested using paired two samples Student’s t-test (ns = P > 0.05, * = P ≤ 0.05, ** = P ≤ 0.01).

This shows that strict N starvation is not applicable for continuously high production levels. Thus, to test the open PGAM valve under varying N availability, exponentially grown seed cultures were washed with BG11^0^ and subsequently cultivated in BG11 containing 5, 10 or 100% nitrate, with the latter corresponding to regular BG11. After 24 hours, the open PGAM valve enabled similar isoprene formation with 5 and 10% nitrate when compared to the control strain cultivated in 100% N, indicating higher carbon flux towards pyruvate in the *pirC* knockout under N-limiting conditions as compared to the wild-type (**Fig. 5F**).

### Sucrose as representative of gluconeogenic routes: Closing the PGAM valve is boosting sucrose production

Sucrose has been shown to accumulate as an osmotic compound to counteract high osmolarity, e.g. in presence of mM concentrations of NaCl, which can be easily tolerated by *Synechocystis* [47,48]. As the PGAM-OFF state increases the carbon flux towards gluconeogenesis, it should enhance sucrose formation upon the addition of salt. First, we tested the impact of salt stress on the strains’ glycogen concentration in both the pgam_KD as well as the pgam_KD pirC_OEX mutant before and after salt shock. The measurement before salt shock was done at 24 h after induction with copper ions, while the measurement after salt shock was done after 120 h of cultivation in salt. Both mutant strains accumulate glycogen during cultivation in medium with 500 mM NaCl. The pgam_KD mutant and pgam_KD pirC_OEX mutant accumulate about ten times more glycogen than the WT (**Fig. 6 A,B**). To monitor sucrose accumulation, both the wild type and the *pgam*_KD strain were subjected to salt shock and different time points after downregulating *pgam* gene expression by copper sulfate were analyzed. For this, both the pgam_KD mutant and the combination mutant of pgam_KD and pirC_OEX were compared to the wild type. All cultures were induced with copper ions 24 h previous to the salt shock. While the pgam_KD mutant did not accumulate more sucrose compared to the WT, the pgam_KD pirC_OEX mutant did accumulate more than twice as much sucrose compared to the WT at 8 h after salt shock (**Fig. 6 C,D**).

**Figure 6:**
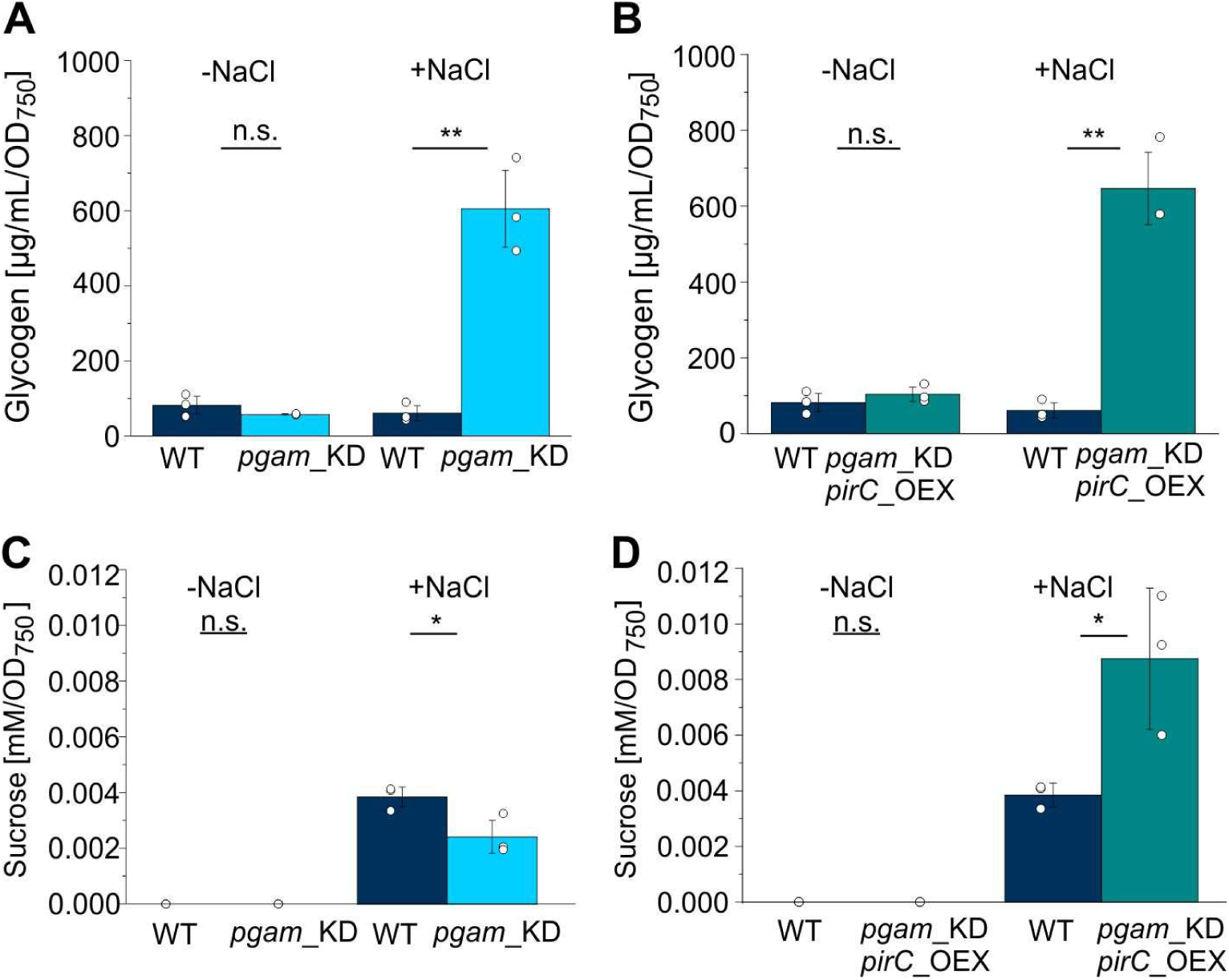
Boosting salt-dependent sucrose formation by modulating the PirC-PGAM switch. For glycogen and sucrose measurements, cells were salt shocked 24 h after induction with copper ions. **A:** Glycogen measurement in the pgam_KD mutant before and after salt shock. **B:** Glycogen measurement in the pgam_KD pirC_OEX mutant before and after salt shock. Glycogen was measured using method B. Measured concentrations are shown with respect to the liquid volume of the sampled cultures at OD750. **C:** Sucrose content in the pgam_KD mutant at two time points before salt shock as well as 8 h after salt shock. **D:** Sucrose content in the pgam_KD pirC_OEX mutant at two time points before and 8 h after salt shock. Each dot represents a biological replicate. T-test was done for statistical analysis (ns = P > 0.05, * = P ≤ 0.05, ** = P ≤ 0.01, *** = P ≤ 0.001).

## Discussion

The 2,3-bisphosphogylcerate-independent phosphoglycerate mutase (PGAM) activity determines whether fixed carbon is retained in gluconeogenic routes supporting glycogen and sucrose synthesis or is directed toward pyruvate, acetyl-CoA, and the TCA cycle. It therefore occupies a strategic position at the interface of the CBB cycle and lower glycolysis and hence, is tightly regulated [22,23]. PGAM activity is controlled by the small protein PirC (also named CfrA in the parallel studies of the Muro-Pastor group, [23]), which inhibits PGAM upon release from PII under nitrogen starvation [22]. Previous work has demonstrated that this key regulatory node has significant influence on cellular carbon allocation and hence is also an attractive target for metabolic engineering [49–52].

In this study, we demonstrate systematic engineering of the PGAM-PirC regulatory node, which represents an effective strategy to redirect central carbon flux in *Synechocystis* toward both native and heterologous products, which is a clear extension to previous studies. By rationally modulating PGAM activity, either by opening or closing the PGAM valve, we established two distinct metabolic states: PGAM-ON and PGAM-OFF, that enable preferential routing of carbon into lower glycolysis or gluconeogenesis, respectively. Importantly, this strategy operates at the level of native metabolic regulation rather than by introducing irreversible pathway blocks, preserving cellular flexibility while enabling productive flux redirection.

To establish the PGAM-ON state, *pgam* was overexpressed using two promoters of different strengths (J23101 and J23119). Under vegetative conditions, increased PGAM abundance did not affect growth, consistent with PGAM reaching equilibrium independently of PirC, which is sequestered by PII. During nitrogen starvation, PirC is released and inhibits PGAM [22], promoting glycogen accumulation. However, PGAM overexpression partially counteracted this inhibition and redirected carbon toward lower glycolysis. As a result, PGAM-ON strains exhibited increased PHB accumulation and a 2-3-fold enhancement in excretion of TCA-cycle-derived metabolites compared to the wild type. Although PGAM overexpression partially counteracts this inhibition, our data indicate that PirC levels are concomitantly upregulated. The reason for this remains elusive. The observed increase in excreted metabolites such as succinate, pyruvate, and 2-oxoglutarate is notable given that wild-type *Synechocystis* releases only trace amounts of organic acids during chlorosis. This highlights the tight endogenous control of metabolite excretion and underscores the effectiveness of PGAM overexpression in overcoming this constraint. The results are consistent with previous studies showing elevated intracellular pools of these metabolites in strains with impaired PirC-mediated regulation [22,33].

Deletion of *pirC* has an even more pronounced effect on carbon allocation. In the Δ*pirC* mutant, PGAM is no longer inhibited during nitrogen starvation, leading to a strong redirection of carbon toward lower glycolysis and the TCA cycle. We show that this metabolic shift not only increases PHB accumulation, as previously reported [49], but also results in substantial excretion of organic acids, particularly 2-oxoglutarate. Given the central role of 2-oxoglutarate in C/N sensing via PII [8], its excretion likely reflects a compensatory response to severe metabolic imbalance. The fact that overexpression of PGAM in the Δ*pirC* background does not further increase metabolite excretion suggests that the system has already reached a new equilibrium, beyond which additional PGAM no longer enhances efflux from the CBB cycle.

Among the excreted metabolites, succinate is of particular biotechnological interest, as it is listed among the top platform chemicals from biomass by the US Department of Energy. *Synechocystis* WT does not excrete succinate under standard photoautotrophic conditions [43], and previous efforts to induce succinate production have relied on deleting carbon sinks such as glycogen synthesis or by disrupting succinate dehydrogenase [53,54]. Our data demonstrate that engineering the PGAM-ON state alone is sufficient to induce succinate excretion, without directly targeting succinate metabolism. This finding positions the PGAM-PirC valve as a complementary strategy that can be combined with established metabolic interventions. For example, coupling the PGAM-ON state with deletion of *sdhB* or introduction of a glyoxylate shunt by expressing an isocitrate lyase from *E. coli* [55] may further enhance succinate yields. In addition, PHB synthesis represents a major acetyl-CoA sink during nitrogen starvation, and its removal could increase carbon availability for organic acid production. Thus, PGAM valve engineering provides a versatile entry point for multi-layered optimization strategies.

Conversely, establishment of a PGAM-OFF state through inducible *pirC* overexpression and/or *pgam* downregulation redirected carbon flux toward gluconeogenesis. All PGAM-OFF strains showed an increase in glycogen accumulation, with glycogen levels peaking approximately 48 h after induction. The strongest effect was observed in our *pirC* overexpression strain, which accumulated glycogen to levels exceeding those previously reported when using analogous genetic configurations based on arsenite-inducible promoters [51]. Beyond glycogen, we explored how the PGAM-OFF state affects sucrose metabolism. Sucrose serves as a compatible solute during salt stress and is metabolically linked to glycogen in *Synechocystis* [48]. When PGAM-OFF strains were subjected to salt shock, both glycogen and sucrose accumulation were enhanced compared to the wild type, although the magnitude of the effect depended on the specific genetic configuration. In particular, the combined *pgam* knockdown/*pirC* overexpression strain exhibited a clear increase in sucrose accumulation, supporting the idea that stronger control over PGAM activity translates into more pronounced flux redirection. These findings align well with recent work demonstrating enhanced sucrose production by controlling carbon flux through CfrA (= PirC) overproduction in *Synechocystis* [52]. Together, these studies reinforce the concept that regulatory control points upstream of sucrose biosynthesis can be leveraged to increase compatible solute production without directly modifying the sucrose synthesis pathway itself. Combining PGAM-OFF engineering with established strategies such as overexpression of the gene of the sucrose phosphate synthase or deletion of the invertase gene [56,57] is therefore a promising route toward further yield improvements.

In addition to native products, we demonstrated that PGAM valve engineering can also improve production of heterologous compounds such as isoprene, derived from pyruvate. Consistent observations have been made in recombinant *Synechocystis* strains optimized for ethanol production [50]. This highlights the broader applicability of this approach for pathways branching from central metabolism. Notably, recent work has identified a PGAM variant with higher activity and reduced sensitivity to PirC inhibition [24], suggesting that future efforts could further decouple PGAM activity from endogenous regulatory feedback and expand the dynamic range of the valve.

## Conclusions

Overall, our study establishes the mechanism controlling the PGAM valve as a powerful tool that can be modulated in *Synechocystis* for controlling carbon flux in a customized manner. By fine-tuning a native regulatory mechanism, we enable flexible redistribution of carbon rather than imposing irreversible pathway blocks that might have further negative effects. More broadly, these findings underscore the potential of small regulatory proteins as engineering targets for metabolic rewiring as suggested previously [19]. The PGAM-PirC valve thus provides a strong foundation for future photosynthesis-driven biotechnological applications and for integrating regulatory engineering with pathway-level optimization.

## Declarations

### Ethics approval and consent to participate

Not applicable

### Consent for publication

Not applicable

### Availability of data and materials

All data generated or analyzed during this study are included in this published article and its supplementary information files.

### Authors’ contributions

SK and KF designed the study. NSB, FH, PB, KO, TO, AK and PF performed research. NSB drafted the manuscript. FH, KF, PB, PL and SK reviewed and edited the manuscript. All authors proof-read the final manuscript and agreed with publication.

## Acknowledgements

We thank Ron Stauder for technical support. ChatGPT was used for language editing of selected manuscript sections, in particular the abstract and parts of the discussion.

