## Supplementary Material for "Conversion of CO_2_ into valuable products: Engineering the PirC-PGAM switch in cyanobacteria to direct carbon flux into desired products" (from another version of this preprint: version 1, 2026-02-08)

### Appendix

#### *Supplementary Methods*

*PGAM/PirC antibody production in rabbits*

The antibodies against PGAM and PirC were obtained from Davids Biotechnologie (Regensburg, Germany) according to the company's protocols. For each protein, three rabbits were immunized after a check of immunity to the cyanobacterial cell extract, performed by Western blot using pre-serum of the rabbits and a cell extract with a whole protein amount of 1 mg (as mentioned previously). Accordingly, rabbits with the lowest reaction against the cell extract were chosen. The immunogenic proteins were produced according to the protein expression protocol described by Orthwein et al. 2025, using the plasmids pET28a-PGAM (PGAM-His) and pJS22 (PirC-His) [24]. To prevent immunization against the His-tags, the tags were removed using the Thrombin CleanCleave™ Kit (Sigma-Aldrich, St. Louis, USA) according to the provided protocol. First, the optimal cleavage times (1 h, 2 h, 4 h, 6 h, and 24 h) were tested. According to the results, 4 h cleavage was used. The thrombin agarose resin was washed twice with 500 µL of 1x cleavage buffer (50 mM Tris-HCl, pH 8.0, 10 mM $CaCl_2$) by resuspension and centrifugation.100 µL of suspended resin was transferred to a 1.5 mL reaction tube and centrifuged. After the buffer was removed, 100 µL of 10x cleavage buffer (500 mM Tris-HCl, pH 8.0, 100 mM $CaCl_2$) was added again. Subsequently, 1 mg of fusion protein was added, and the slurry was diluted to 1 ml to obtain the cleavage slurry. The cleavage was done by gentle agitation within a reaction tube mixer for 4 h. The resin was removed by gentle centrifugation, and the supernatant was transferred to a new reaction tube. The buffer was exchanged to the transport buffer (50 mM Tris-HCl, 50 mM NaCl). This procedure was done with 5 vials. After cleavage, the proteins were pooled to a final concentration of 1 mg/mL in a transport buffer. Three vials of each protein were sent to David's Biotechnology in a cooling box (~4 °C) for immunization. The final serums were tested via Western blot according to the protocol (as mentioned previously).

#### *Supplementary Tables*

**Table S1: List of plasmids used and generated in this study.**

| Name | Plasmid | Purpose | Source |
|---|---|---|---|
| | pSEVA251, $Km^R$ | Control plasmid | [58] |
| | pSEVA451, $Sm^R$ | Control plasmid | [58] |
| pNB204 | pSEVA251-$P_{J23101}$::*pgam*, $Km^R$ | Overexpression of *slr1945* with J23101 promoter | This study |
| pNB205 | pSEVA251-$P_{J23119}$::*pgam*, $Km^R$ | Overexpression of *slr1945* with J23119 promoter | This study |
| pFH01 | pSEVA451-$P_{petE}$::*pirC, $Spec^R$* | Overexpression of *sll0944* with the promoter of the *petE* gene | This study |
| pFH10 | pEX-K248-$P_{petJ}$::*pgam, $Km^R$* | Downregulation of *slr1945* with the promoter of the *petJ* gene | This study |
| pEERM3:2MEP-IspS | pJ344, $Km^R$ | Knockout of *slr0168* replacing it with Ptrc-driven overexpression of DXS, Idi, IspS and Kanamycin resistance casette | [46] |
| pEERM3-PirC:2MEP-IspS | pJ344, $Km^R$ | Knockout of *sll0944* replacing it with Ptrc-driven overexpression of DXS, Idi, IspS and Kanamycin resistance casette | This study |
| pJS22 | pET15b, $Amp^R$ | Overexpression of His8-PirC with the T7 promoter | [22] |
| pET28a_PGAM | pET28a, $Km^R$ | Overexpression of PGAM-His6 with the T7 promoter | [22] |

**Table S2: List of strains used and generated in this study.**

| Organism/ Strain | | Genotype | Purpose/ Reference |
|---|---|---|---|
| *E. coli* | NEB10β | Δ(ara-leu) 7697 araD139 fhuA ΔlacX74 galK16 galE15 e14- Φ80dlacZΔM15 recA1 relA1 endA1 nupG rpsL (StrR) rph spoT1 Δ(mrr-hsdRMS-mcrBC) | Molecular Cloning, NEB |
| | Stellar | F- endA1 supE44 thi-1 recA1 relA1 gyrA96 phoA Φ80dlacZΔM15 Δ(lacZYA-argF) U169 Δ(mrr-hsdRMS-mcrBC), ΔmcrA, λ- | Molecular Cloning, TAKARA |
| | Top10 | $F^{-}$*mcr*A Δ(*mrr-hsd*RMS-*mcr*BC) φ80*lac*ZΔM15 Δ*lac*X74 *rec*A1 *ara*D139 Δ(*ara-leu*)7697 *gal*U *gal*K $\lambda^{-}$*rps*L($Str^{R}$) *end*A1 *nup*G | Molecular cloning, |
| | DH5α | $F^{-}$ φ80*lac*ZΔM15 Δ(*lac*ZYA-*arg*F)U169 *rec*A1 *end*A1 *hsd*R17($r_K^-$, $m_K^+$) *pho*A *sup*E44 $\lambda^{-}$*thi*-1 *gyr*A96 *rel*A1 | Molecular cloning, |
| *Synechocystis* sp. PCC 6803 | wildtype glucose-tolerant | WT | Control strain |
| | Δ*pirC* | *sll0944*::$Spec^R$ | [22] |
| | 101_*pgam* | WT + pSEVA251-$P_{J23101}$::*pgam*, $Km^R$ | This study |
| | 119_*pgam* | WT + pSEVA251-$P_{J23091}$::*pgam*, $Km^R$ | This study |
| | Δ*pirC* + 101_*pgam* | *sll0944*::$Spec^R$ + pSEVA251-$P_{J23101}$::*pgam*, $Km^R$ | This study |
| | pirC_OEX | WT + pSEVA451-$P_{petE}$::*pirC*, $Sm^R$ | This study |
| | pgam_KD | $P_{petJ}$::pgam, $Km^R$ | This study |
| | NSI-IspS | *slr0168:dxs-idi-ispS::*$Km^R$ | [46] |
| | ΔPirC-IspS | *sll0944:dxs-idi-ispS::*$Km^R$ | This study |

**Table S3: List of oligonucleotides used in this study.**

GA= Gibson Assembly, sdm= side-directed mutagenesis, AQ = AQUA cloning

| name | sequence | Purpose, Reference |
|---|---|---|
| NB042 | aaaagcgcggatcctctagagtcgacctgca | GA, pSEVA-$P_{J23101}$::*pgam* (This study) |
| NB045 | gtcgactctagaggatccgcgcttttc | |
| NB046 | ctgtaaa*gaattcgcgcggcc* | |
| NB047 | ccgcggccgcgcgaattctttacagctagctcagtcc | |
| NB062 | agggcggcggatttgtcc | Colony PCR (https://seva-plasmids.com/backbone-modules-nomenclature/) |
| NB063 | gcggcaaccgagcgttc | |

| | | |
|---|---|---|
| NB075 | cctaggtataatgctagcATCTTTTGCAGACTTTATAACTG TAGTGAG | sdm, substitute $P_{J23101}$ with $P_{J23119}$ on pSEVA251-$P_{J23101}$::*pgam* plasmid (This study) |
| NB076 | actgagctagctgtcaaGAATTCGCGCGGCCG | |
| FH013 | gaattcGAAGGGATAGCAAGCTAATTTTT | Ligation for pSEVA451-$P_{petE}$::*pirC* (This study) |
| FH017 | tttgggacatACTTCTTGGCGATTGTATCTAT | |
| FH018 | gccaagaagtATGTCCCAAATTCTTGATCCC | |
| FH019 | ggatccTTTTTCTTAGTGCTTTGTCCG | |
| FH020 | AGGGAGTCCCGGATTAAA | Colony PCR pgam_KD (This study) |
| FH027 | GCGCTTTTCCATCCCC | |
| FH112 | ATCTTTTGCAGACTTTATAACTGTAG | |
| 2-MEP-PirCup_fw | aatcaccaacgccaaaAAAAAAAGGATCTCAAGAAGAT CCTT | AQ, pEERM3-PirC:2MEP-IspS (This study) |
| 2-MEP-PirCdwn_rev | ttcattaattcattgGGATCCCTCGAGTCCCG | |
| PirCup_fw | CCTATAGTGTCTTCGGGGattcgccgtaataggcactg | |
| PirCup_2MEP _rev | TTGAGATCCTTTTTTTtttggcgttggtgattactt | |
| PirCdwn-2MEP_fw | GGGACTCGAGGGATCCcaatgaattaatgaattggaatac | |
| PirCdwn-rev | GACGTATATGGTCTTCTTTTatctccgccatagatcctt | |
| pEERM-PirCdwn_fw | gaaggatctatggcggagatAAAAGAAGACCATATACGTC | |
| pEERM-PirCup_rev | cagtgcctattacggcgaatCCCCGAAGACACTATAGG | |

#### *Supplementary Figures*

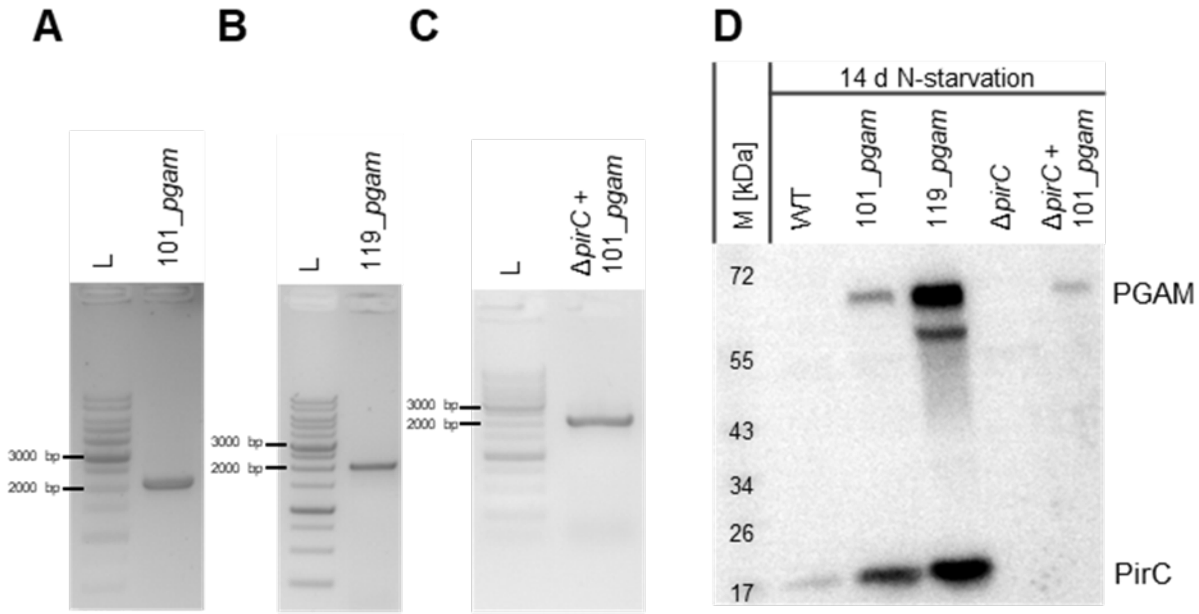


**Suppl. Figure 1: Colony PCR of strains 101_*pgam*, 119_*pgam* and Δ*pirC* + 101_*pgam and* immunoblots against PGAM and PirC. A-C:** Colony PCR of the *Synechocystis strains* 101_*pgam* (A), 119_*pgam* (B) and Δ*pirC* + 101_pgam (C) with the primers NB062/063. As ladder (L) the GenLadder 1kb ready-to-use DNA-Marker from Genaxxon was used. **D:** Immunoblots against PGAM and PirC 14 days into nitrogen starvation.

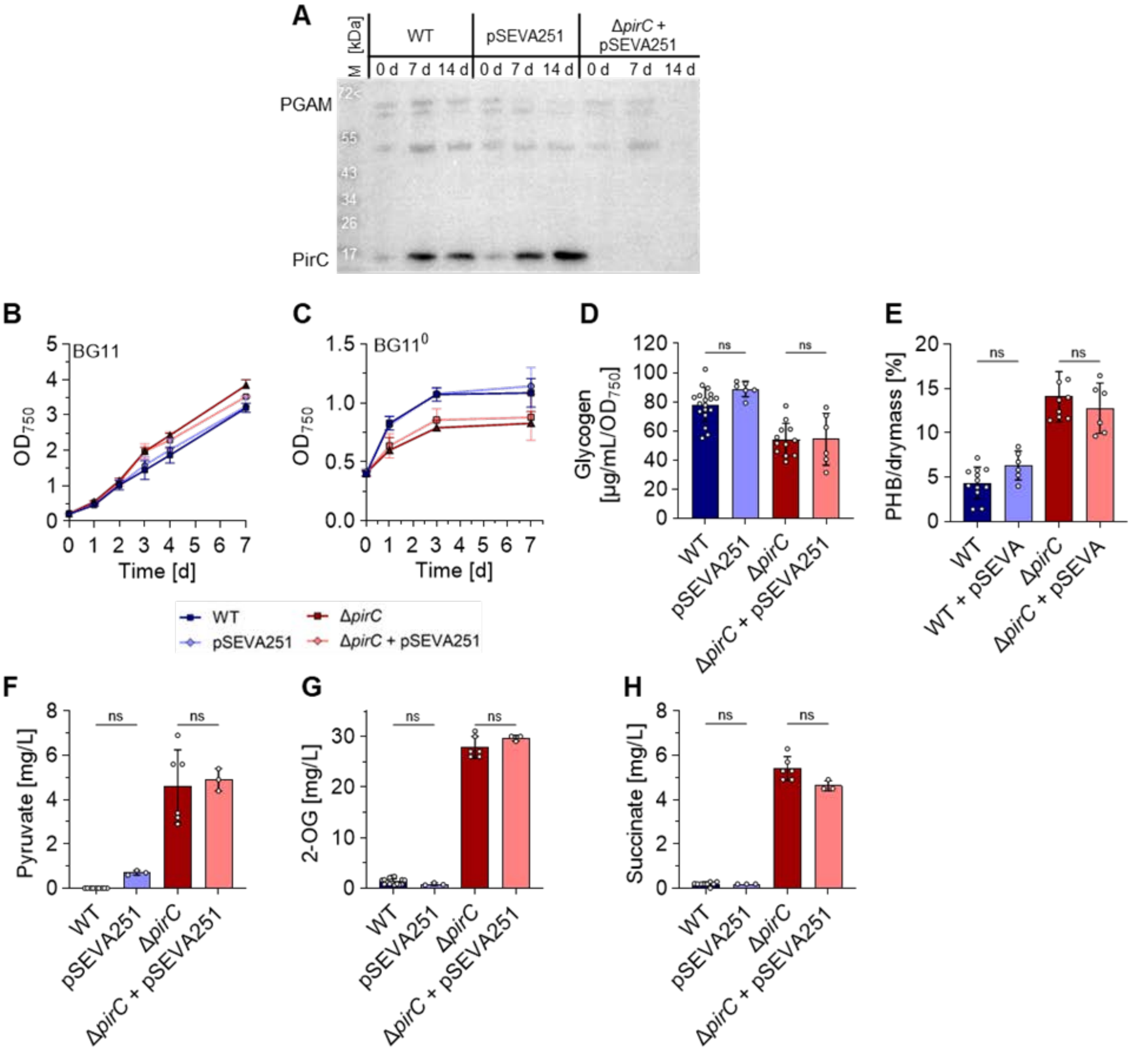


**Figure S2: Empty pSEVA251 plasmid has no effect on growth, glycogen and PHB level. A:** Immunoblots against PGAM and PirC in vegetative conditions (0 d) and 7 days into nitrogen starvation (7 d). **B-C:** Vegetative growth (B) and $OD_{750}$ under nitrogen starvation (C) with at least three biological replicates. **D-E:** Quantification of glycogen with method A (D) and PHB (E) 7 days in nitrogen starvation. **F-I:** Extracellular metabolites were quantified with HPLC-MS from the supernatant of 7-day nitrogen-depleted cell cultures. Each dot represents a biological replicate. Dunn's multiple comparisons test was used (ns = P > 0.05, * = P ≤ 0.05, ** = P ≤ 0.01, *** = P≤ 0.001).
